# MAPS integrates overlooked regulation of actin-targeting effector SteC into the virulence control network of *Salmonella* small RNA PinT

**DOI:** 10.1101/2020.07.28.223677

**Authors:** Sara Correia Santos, Thorsten Bischler, Alexander J. Westermann, Jörg Vogel

## Abstract

A full understanding of the contribution of small RNAs (sRNAs) to bacterial virulence demands knowledge of their target suites under infection-relevant conditions. Here, we take an integrative approach to capturing targets of the Hfq-associated sRNA PinT, a known post-transcriptional timer of the two major virulence programs of *Salmonella enterica*. Using MS2 affinity purification and RNA-sequencing (MAPS), we identify PinT ligands in bacteria under *in-vitro* conditions mimicking specific stages of the infection cycle, and in bacteria growing inside macrophages. This reveals PinT-mediated translational inhibition of the secreted effector kinase SteC, which had gone unnoticed in previous target searches. Using genetic, biochemical, and microscopic assays, we provide evidence for PinT-mediated repression of *steC* mRNA, eventually preventing actin rearrangements in infected host cells. Our findings support the role of PinT as a central post-transcriptional regulator in *Salmonella* virulence and illustrate the need for complementary methods to reveal the full target suites of sRNAs.

## INTRODUCTION

To successfully initiate and sustain an infection, bacterial pathogens possess complex regulatory networks enabling them to precisely time the synthesis of their virulence proteins. Timing is crucial: if expressed too early, virulence factors and their export machineries add a substantial metabolic cost and the risk of premature sensing of a pathogen by the host. If expressed too late, the pathogen might fail to establish its protective niche in time, risking clearance by host defense mechanisms. Much of this control takes place at the DNA level, and responsible transcriptional regulators are now known for many pathogens (Cabezas et al., 2018; Colgan et al., 2016; Ellermeier and Slauch, 2007; Pérez-Morales et al., 2017; Smith et al., 2016). Starting with pioneering work on RNAIII in *Staphylococcus aureus*, bacteria have also increasingly been shown to use regulatory RNAs to integrate virulence factor production with quorum sensing, biofilm formation, and nutrient status (Guillet et al., 2013). Nonetheless, although bacterial pathogens have been shown to express many small RNAs (sRNAs) during infection (Westermann, 2018), the roles of noncoding RNA in timing virulence programs remain little understood.

In Gram-negative pathogens, evidence for sRNA control of virulence has been twofold. First, genetic inactivation of the two major sRNA-binding proteins that facilitate sRNA-mediated regulation of mRNAs, Hfq and ProQ, typically attenuates infectivity of many species (Ansong et al., 2009; Chao and Vogel, 2010; Sittka et al., 2007; Westermann et al., 2019). Second, over the years, several mRNAs of virulence-associated proteins were identified as sRNA targets in diverse Gram-negative pathogens (Bradley et al., 2011; Gong et al., 2011; Murphy and Payne, 2007; Padalon-Brauch et al., 2008; Sievers et al., 2014), including transcripts related to virulence-associated processes such as quorum sensing and biofilm formation (Bardill et al., 2011; Lenz et al., 2004; Papenfort et al., 2015; Shao and Bassler, 2014; Sonnleitner et al., 2011).

Working in the model species *Salmonella enterica* serovar Typhimurium (henceforth, *Salmonella*), we recently identified a ~80-nt sRNA called PinT, which similarly to *S. aureus* RNAIII, seems to play a central role in timing virulence factor expression of this intracellular pathogen (Westermann et al., 2016). PinT is the top-induced noncoding transcript after *Salmonella* enters eukaryotic host cells (Westermann et al., 2016). Controlled by the PhoP/Q two-component system, this sRNA is co-activated with the physically unlinked *Salmonella* pathogenicity island 2 (SPI-2) that encodes a type-III secretion system (T3SS) apparatus and corresponding effector proteins required for intracellular survival.

Three major functions of PinT have been established. Firstly, by Hfq-dependent seed pairing, PinT downregulates specific virulence factor mRNAs (*sopE, sopE2*) from the SPI-1 invasion gene program (Westermann et al., 2016). Secondly, PinT also inhibits invasion gene expression globally, by repressing the mRNAs of two major SPI-1 transcription factors, HilA and RtsA (Kim et al., 2019). Thirdly, while hastening the shutoff of SPI-1, PinT delays full activation of SPI-2, by inhibiting the synthesis of the general transcription factor CRP and of the SPI-2-encoded transcriptional regulator SsrB (Kim et al., 2019; Westermann et al., 2016). In other words, PinT acts as a post-transcriptional timer, shaping the transition from one (SPI-1, invasion) to the other (SPI-2, intracellular lifestyle) major virulence programs of *Salmonella*. The combined activities of PinT in *Salmonella* have a pervasive molecular effect on host cells, with ~10% of all host mRNAs showing altered expression when infected with Δ*pinT* versus wild-type bacteria (Westermann et al., 2016). Yet, PinT shares with many other sRNAs the limitation that a knockout produces no robust macroscopic phenotype in standard cell culture or mouse models (Barquist et al., 2016). Therefore, to understand the full scope of PinT activity, a comprehensive analysis of its mRNA targets is needed.

Thus far, PinT targets have been predicted by pulse-overexpression of the sRNA and scoring global changes in mRNA levels (Westermann et al., 2016), or by educated guesses combined with *in-silico* predictions of base complementarity (Kim et al., 2019). These routes have clear limitations, e.g., sRNA pulse-expression misses mRNA targets regulated only on the level of translation, but not transcript stability. Therefore, for a comprehensive view of PinT targets and mechanisms, we here apply an orthogonal approach called MAPS (Lalaouna et al., 2015), which captures physical sRNA-RNA interactions in bacterial cells. In MAPS, an sRNA of interest is typically fused to an MS2 aptamer to enable recovery from cell lysates by affinity chromatography (Said et al., 2009), followed by RNA-seq analysis of co-purifying transcripts. Originally developed for the iron-responsive RyhB sRNA in *E. coli* (Lalaouna et al., 2015), MAPS has since uncovered many unrecognized targets of other well-characterized *E. coli* sRNAs (Lalaouna et al., 2018, 2019b) and enabled global target screens in other bacterial species (Georg et al., 2020; Lalaouna et al., 2019a; Silva et al., 2019; Tien et al., 2018; Tomasini et al., 2017).

While these previous studies often monitored a single experimental condition and overexpressed the sRNA of interest to high levels, we here take an integrative MAPS approach under *bona fide* infection conditions, including bacterial growth inside macrophages, and seek to capture PinT targets at physiological concentrations of the sRNA (Fig. 1a). This multi-condition analysis predicts a previously unrecognized PinT-mediated translational repression of the secreted SPI-2 effector SteC. Physiologically, this adds regulation of effector-induced actin rearrangement in epithelial cells to the intracellular activities of PinT. The integrative MAPS approach reported here should facilitate bottom-up analysis of sRNA targets during the intracellular phase of other bacterial pathogens.

**Figure 1.**
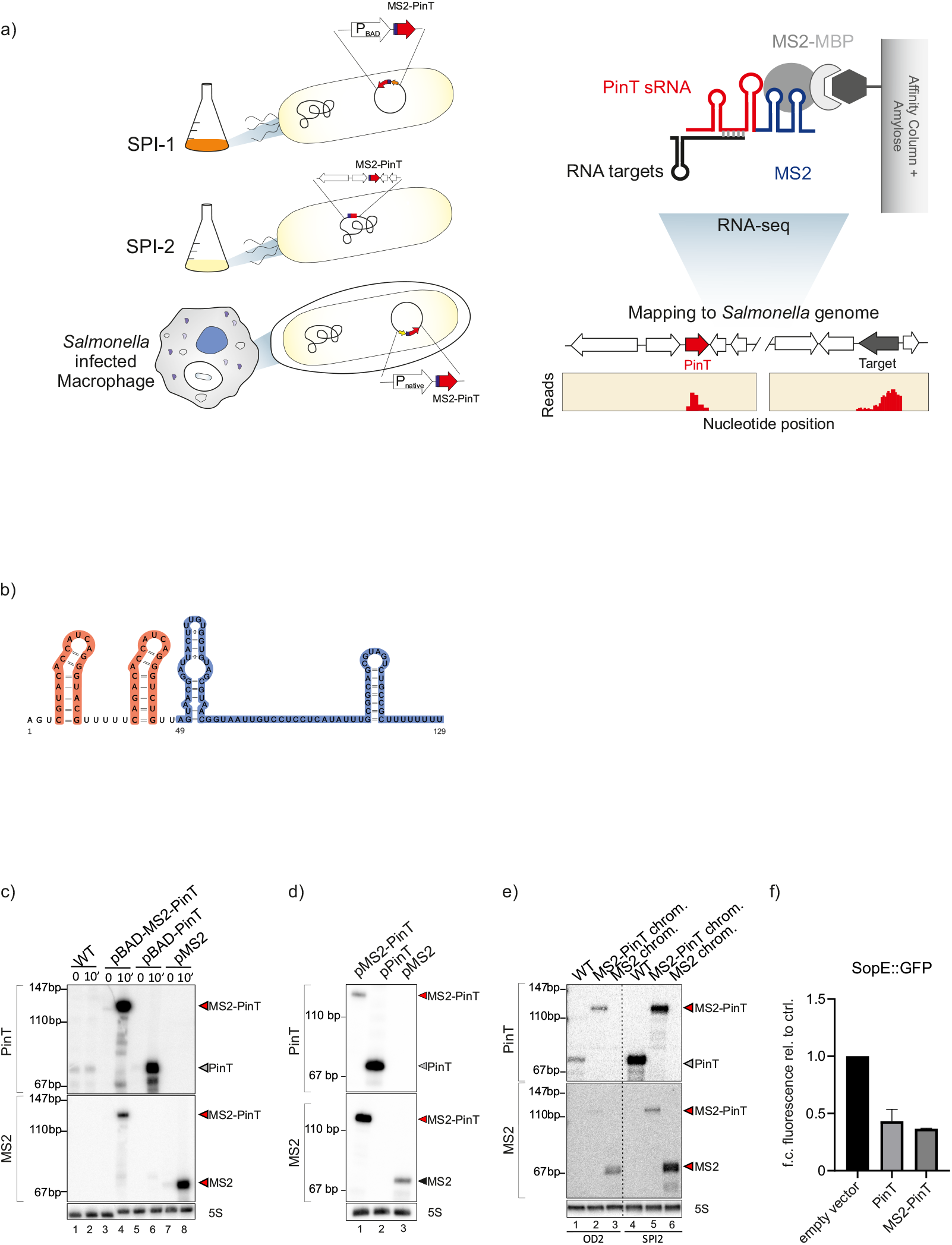
**a)** Experimental flow of MAPS in *Salmonella*. **b)** Secondary structure prediction of MS2-PinT using mfold web server and VARNA applet for visualization of RNA secondary structure. **c-e)** Northern Blot analysis of **c)** *Salmonella* Typhimurium SL1344 wild-type carrying empty vector control (lanes 1-2 - pKP8-35), or the *pinT* deletion strain carrying a plasmid expressing the wild-type PinT (lanes 5-6 - pYC5-34); the aptamer tagged PinT, i.e. MS2-PinT (lanes 3-4 – pYC310); or the tags alone, MS2 (lanes 7-8 – pYC310) under an arabinose inducible promoter; before or after 10 minutes of induction; **d)** *Salmonella* Typhimurium SL1344 wild-type carrying empty vector control (lane1 - pJV300), or the *pinT* deletion strain carrying a plasmid constitutively expressing either the wild-type PinT (lane 2 – pYC55), the MS2-PinT (lane 3 – pSS31) or the MS2 aptamer (lane 4 – pSS32), under the control of the PinT native promoter; **e)** of *Salmonella* wild-type (lanes 1,4) or *Salmonella* carrying a chromosomal copy of either MS2-PinT (lanes 2, 5) or of MS2 (lanes 3, 6); grown under SPI-1 or SPI-2 inducing conditions. Includes 5S as a loading control. **f)** GFP reporter assay. *sopE::gfp* reporter gene fusion (including the complete 5’UTR and the first 60 codons) was used to measure of the interaction between the pulse expressed wild-type (PinT) or the tagged PinT (MS2-PinT). Results correspond to the representation of three biological replicates.

## RESULTS

### Establishing MAPS for Salmonella PinT sRNA

MAPS requires an sRNA to tolerate fusion to a relatively large aptamer without compromising its intracellular stability, RBP association (if applicable) or base pairing activity (Lalaouna et al., 2017). Adding a 48-nt MS2 aptamer to the 5’ end of PinT (~80 nt) generates a 129-nt RNA fusion with no predicted distortion of the folding of the linked PinT (Fig. 1b). To validate its functionality in *Salmonella*, we first expressed the MS2-PinT fusion from an arabinose-inducible, plasmid-borne promoter. Following induction for 10 min in early stationary phase (OD_600nm_ of 2.0), the MS2-PinT construct yielded the same amount of sRNA compared to an analogous expression vector harboring wild-type PinT (Fig. 1c). Importantly, the MS2-PinT sRNA accumulated as a single species, i.e. the aptamer did not cause aberrant processing. Similar results were obtained with the MS2 sequence directly inserted into a plasmid-borne *pinT* gene under its native promoter, proving that the insertion did not compromise endogenous transcription of *pinT* (Fig.1d). Finally, we evaluated whether the MS2-PinT sRNA was functional, testing its ability to repress the well-characterized *sopE* mRNA target (Westermann et al., 2016). Using a *sopE::gfp* gene fusion as readout, we observed equal repression by wild-type PinT and the MS2 fusion (Fig. 1f). Together, these experiments showed that the aptamer impacted neither Hfq association nor target pairing, qualifying the MS2-PinT fusion for MAPS-based target capture in *Salmonella*.

### MAPS recapitulates major PinT targets and identifies new candidates

To establish MAPS-based target capture for PinT, we first performed the original MAPS protocol in *in-vitro* cultures of *Salmonella*. Three different plasmids were used: the control plasmids pBAD-PinT and pBAD-MS2, expressing the untagged sRNA or the aptamer alone, respectively; and pBAD-MS2-PinT carrying the arabinose-inducible fusion sRNA. We induced these plasmids in LB cultures grown to an OD_600nm_ of 2.0, which is a condition when SPI-1 is activated while the endogenous PinT sRNA exhibits intermediate expression (Westermann et al., 2016). Because overexpressed PinT was known to rapidly induce target mRNA degradation, we optimized the induction time (Supplementary Fig. 1a). We settled on 2 min, when PinT is already abundant and the *sopE* and *sopE2* mRNAs—two well-characterized targets (Westermann et al., 2016)—start decaying.

To rank putative targets recovered with the MS2-PinT sRNA from *Salmonella* lysates, we calculated fold enrichment comparing normalized read counts from RNA-seq of the MS2-PinT and the untagged PinT samples, each collected in duplicate (Fig. 2a, Supplementary Table 1). Known PinT targets (*sopE, sopE2, grxA, crp, hilA, rtsA and ssrB*) were enriched in the MS2-PinT pull-down, compared with PinT alone, confirming that the fusion sRNA was functional (Fig. 2a). Surprisingly, one of the most enriched transcripts was *steC* (rank#4; Supplementary Table 1), a virulence factor encoding mRNA that had gone unnoticed in all previous PinT target searches, and to which we will return further below.

**Figure 2.**
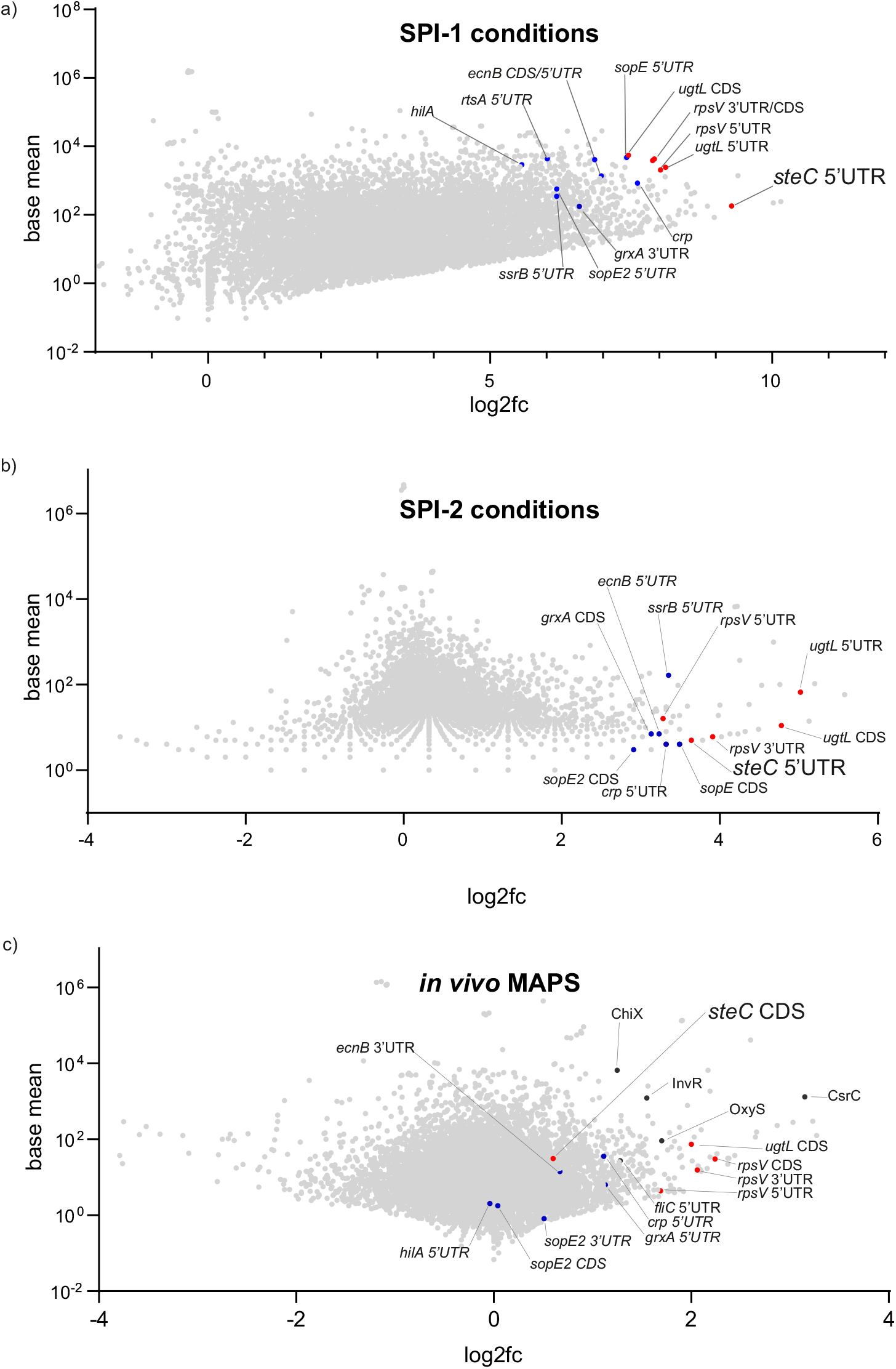
MAPS pull downs in **a)** SPI-1 conditions; **b)** SPI-2 conditions and **c)** *in-vivo* MAPS. MA plots show log_2_ fold change in enrichment in MS2-PinT relative to PinT. Data were collected from two biological replicates. New candidate targets are labeled in red and previously known targets in blue. Candidate sRNA targets are labeled in black. All other transcripts are labeled in light grey.

### MAPS at physiological sRNA concentrations, inside host cells

All MAPS studies so far have relied upon overexpressed MS2 fusions to reach sufficiently high levels for target pull-down. However, we reasoned that the high abundance of native PinT under infection conditions (Westermann et al., 2016) would allow for MAPS under native conditions. To test this, we performed MAPS with a *Salmonella* strain in which the MS2 tag had been engineered into the chromosomal *pinT* locus (Fig. 1a). Grown in a minimal medium that mimics the intracellular environment of host cells (Löber et al., 2006), this strain produced comparable levels of PinT as the wild-type strain (Fig. 1e). MAPS under this condition again captured previously described targets of PinT as well as the above identified candidate *steC* (Fig. 2b, Supplementary Table 1).

To identify PinT targets in a true infection setting, we performed MAPS after host cell invasion, on bacteria replicating inside macrophages. In this setting, to guarantee robust sRNA recovery, we introduced plasmids expressing MS2-PinT, untagged PinT, or the MS2 tag from the native *pinT* promotor, in a *pinT* deletion strain (Fig. 1a). Using a multiplicity of infection (MOI) of 50, 6% of macrophages contained intracellular *Salmonella*. At 4 h after infection, macrophages were lysed and the recovered bacteria subjected to MAPS. Of the ~30 million cDNA reads obtained per library, 5-46% mapped to the eukaryotic host transcriptome. The majority of the *Salmonella*-specific reads were from rRNA, leaving 3-18% of other bacterial reads for the identification of PinT targets.

Intracellular targets of PinT were predicted by calculating enrichment in MS2-PinT over the untagged PinT control (Fig. 2c, Supplementary Table 1). Of known targets, *grxA* and *crp* were slightly enriched with the MS2-PinT (log_2_ f.c of 1.13 or 1.11, respectively). The remaining targets, *sopE, sopE2, hilA, rtsA* and *ssrB*, are underrepresented in the MS2-PinT pull down samples (log_2_ f.c ≤1), as expected since these genes are expressed earlier during infection. In other words, MAPS at this 4h post-infection (p.i.) time point *in-vivo* differs from the *in-vitro* SPI-2 condition. Other enriched mRNAs of interest included *ugtL* encoding a membrane protein involved in PhoQ activation, *rpsV* encoding the 30S ribosomal subunit protein S22 and *fliC*, encoding for the major flagellin (Fig. 2c, Supplementary Table 1). As with bacteria from SPI-2 media, this MAPS experiment on intracellular bacteria again recovered the *steC* mRNA, although the enrichment was not as strong as under pre-infection conditions (log_2_ f.c. 0.60 inside macrophages vs. log_2_ f.c. 9.28 at OD_600nm_ of 2.0).

### Pin Tregulates steC on the post-transcriptional level

Coincidentally, we had previously used the *steC* promoter as a readout for PinT activity on CRP, assuming that this sRNA regulated the *steC* gene indirectly as part of its global effect on SPI-2 activation (Westermann et al., 2016). However, the integrated MAPS data now suggested that PinT also regulates *steC* directly, through an RNA-RNA interaction. SteC is an effector kinase secreted through the T3SS of SPI-2 into the host cytosol, where it induces the assembly of an F-actin meshwork around the *Salmonella* containing vacuole (SCV), thereby restraining bacterial growth (Odendall et al., 2012; Poh et al., 2008). Therefore, in the following we focused on the characterization of PinT-mediated SteC regulation and its impact on host cells during infection. To test the predicted direct regulation by PinT, we determined changes of *steC* mRNA levels after induced expression of PinT (Fig. 3a). We observed a rapid decrease in *steC* mRNA levels, disappearing at roughly the same rate as the established direct target, *sopE* (Fig. 3a). In addition, we constructed a *Salmonella* strain in which we fused a FLAG epitope to the C-terminus of the *steC* reading frame, for western blot detection of the protein. Using this *steC*::3xFLAG strain, we observed that while *pinT* deletion mildly increased protein levels, constitutive PinT expression fully depleted the SteC protein (Fig. 3b).

**Figure 3.**
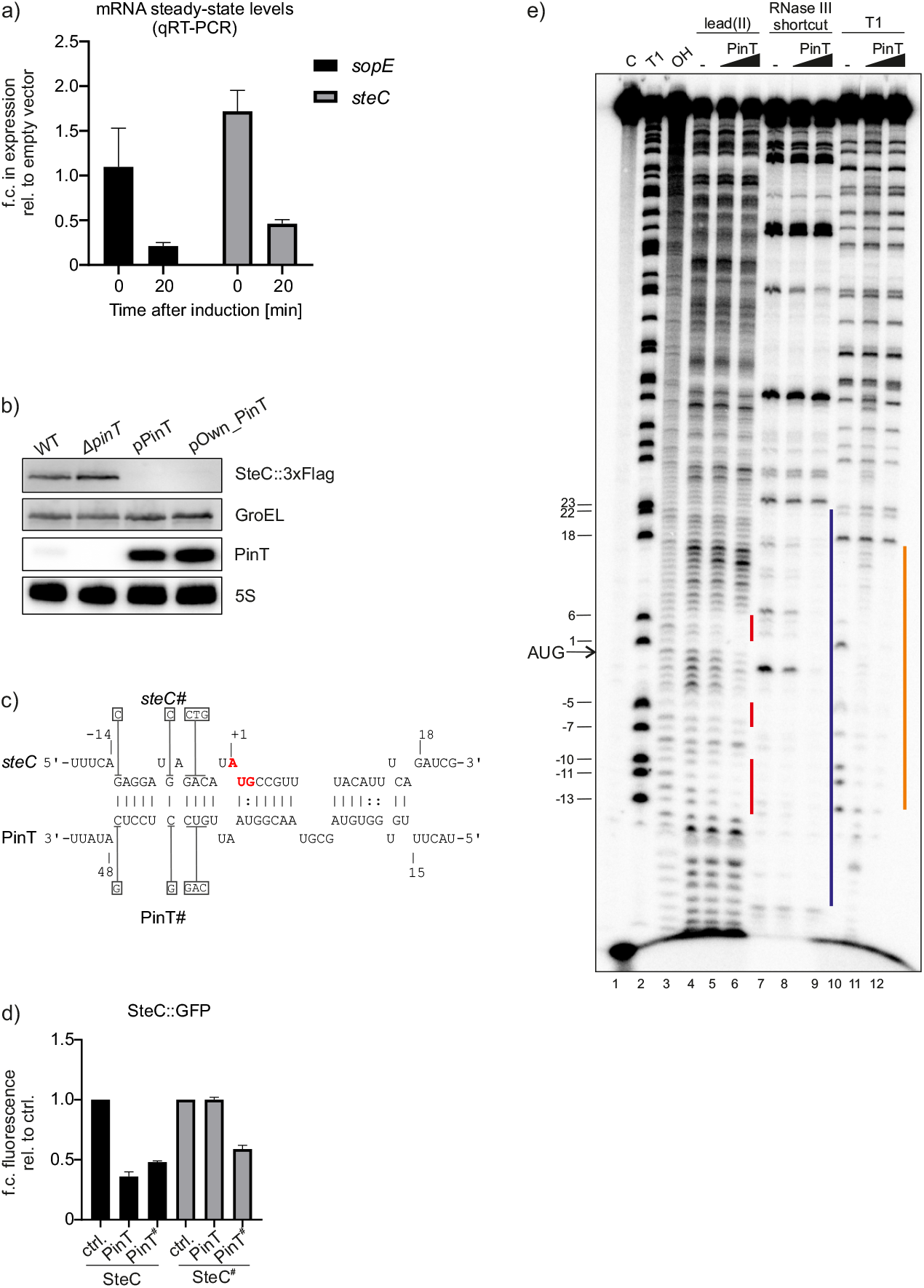
**a)** qRT-PCR measurements of *sopE* and *steC* mRNAs before and after 20 minutes of pulse expression of PinT. Transcript fold change expression was compared with empty vector before and after induction. Results correspond to the representation of three biological replicates. **b)** Northern blot and Western-blot analysis of SteC::3xFlag in a *pinT* deletion strain, or a complemented strain where PinT expressed from a plasmid under the control of a constitutive promotor or of its own promotor. Includes GroEL and 5S as a loading control for protein and RNA, respectively. **c)** *in-silico* prediction of interaction between *steC* mRNA and PinT. PinT# and *steC#* mutations are show in boxes; the start codon of *steC* is highlighted in red. **d)** GFP reporter assay. *steC::gfp* or *steC#::gfp* reporter gene fusion was used to measure of the interaction between the pulse expressed wild-type (PinT) and mRNA fusion, on a *pinT* deletion background. All results correspond to the representation of three biological replicates. **e)** *in-vitro* structure probing for identification of PinT binding sites on *steC* mRNA truncated variant (from TSS to +250nt in the CDS). 5’-End-labeled *stec* mRNA (5 nM) was subjected to RNase T1, lead(II), and Shortcut RNaseIII cleavage in the absence (lanes 4, 7, 10) and presence of different concentrations of unlabelled PinT RNAs (0.5 μM final concentration in lanes 5, 8, 9; 5 μM final concentration in lanes 6, 9, 12). (Lane C) Untreated GcvB RNA. (Lane T1) RNase T1 ladder of hydrolyzed denatured *steC*. (Lane OH) Alkaline ladder. Vertical bars indicate the *steC* region protected by PinT.

Hfq-associated sRNAs such as PinT typically repress mRNAs by antisense sequestration of the translation initiation region (Hör et al., 2020). Using *in-silico* analysis for base pairing regions, we predicted that PinT uses its previously determined seed sequence (Westermann et al., 2016) to form a ~17 bp duplex with the Shine Dalgarno (SD) sequence and AUG start codon of *steC* (Fig. 3c, Supplementary Fig. 2a). Testing the presumed inhibition, we found that PinT indeed repressed a *steC::gfp* translational fusion, which includes the 30 nucleotide-long 5’ UTR and the first 21 codons of *steC* (Fig. 3d). Importantly, since this fusion is transcribed from a constitutive promoter, repression must occur post-transcriptionally (Fig. 3d).

Using *in-vitro* synthesized PinT sRNA and a 5’ fragment of *steC* mRNA, we observed that they readily formed a complex when incubated with each other, as shown by the electrophoretic mobility shift assay (EMSA) (Supplementary Fig. 2b-c). Binding occurred with an apparent dissociation constant of 533 nM (Supplementary Fig. 2e), similar to other established sRNA-mRNA interactions without addition of Hfq (Bobrovskyy et al., 2019). When Hfq was added to the binding reaction, a further shift occurred, predicting the formation of a super complex of the two RNA partners together with Hfq (Supplementary Fig. 2d-e).

Next, we used structure probing to experimentally define the putative PinT-*steC* RNA duplex. A fixed amount of an *in-vitro*-transcribed, radiolabeled segment of *steC* mRNA (encompassing the 5’UTR and the first 21 codons) was treated with lead (II)-acetate, RNase T1 or shortcut RNase III, without or with increasing concentrations of PinT. Visualization of the resulting cleavage products in a denaturing gel supported the *in-silico* prediction that the *steC* start codon is sequestered as a result of pairing with PinT (Fig. 3e).

The *in-vitro* probing results allowed us to select critical bases to mutate in PinT and *steC*, finally proving a direct RNA interaction *in-vivo*. Specifically, we generated a *steC#::gfp* fusion with five bases mutated in the region upstream of the start codon (Fig. 3c); these changes fully abrogated repression by PinT (Fig. 3d). However, compensatory changes in the sRNA (PinT# variant) fully restored repression of the *steC#::gfp* fusion. Conversely, PinT# also repressed the wild-type *steC::gfp* reporter, albeit less strongly than did wild-type PinT, maybe as a combined effect of PinT# overexpression and the potential for seven nucleotides to form base pairs even in the mutated sRNA.

### Evidence for translational repression of SteC by PinT

The location of the PinT target site within *steC* 5’UTR pointed to a direct inhibition of translation initiation. We obtained evidence for this mechanism in two different types of *in-vitro* assays. Firstly, we translated the *steC::*3xFLAG mRNA using 70S ribosomes, in the absence or presence of PinT or Hfq (Fig. 4a). We observed clear inhibition of SteC::3xFLAG protein synthesis by increasing concentrations of PinT, provided Hfq was added to the reaction (Fig. 4a, lanes 9-10). By contrast, PinT or Hfq alone did not inhibit *steC::*3xFLAG mRNA translation (lanes 3-4 and 7). As expected, translation of the unrelated *hupA::*3xFLAG mRNA, used as a negative control, was unaltered by PinT or Hfq (lanes 5-6, 8 and 11-12).

**Figure 4.**
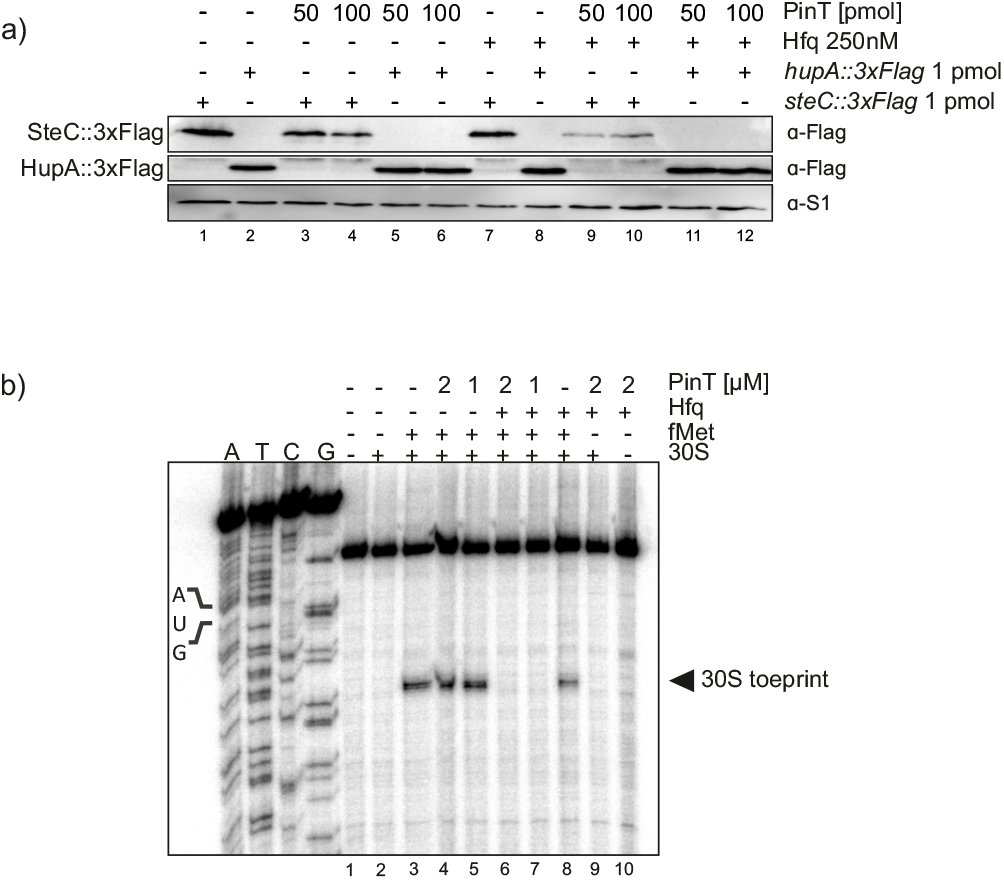
**a)** *in-vitro* translation assay, full-length *steC::3xFLAG or hupA::3xFLAG* mRNA fusion were *in-vitro* translated, individually, with reconstituted 70S ribosomes in the presence or absence of PinT and/or Hfq. **b)** Toeprinting assay on the *steC::gfp* mRNA performed in the presence or absence of PinT and/or Hfq.

Secondly, to prove that PinT inhibited the initiation step of translation, we performed 30S ribosome toeprinting assays (Hartz et al., 1988). The *steC::gfp* mRNA was annealed to an end-labeled primer, complementary to the *gfp* coding region, and incubated with 30S subunits in the presence or absence of uncharged tRNA^fMet^, followed by reverse transcription. Analysis of the extension products revealed one ribosome-induced, tRNA^fMet^-dependent toeprint at the characteristic +15 nt position (Fig. 4b, lane 3). This toeprint signal strongly decreased in the presence of both PinT and Hfq (lanes 6, 7). Addition of Hfq alone, which has been reported to suffice for repression of some mRNAs (Chen and Gottesman, 2017; Ellis et al., 2017; Sonnleitner and Bläsi, 2014), did not affect the *steC* toeprint (lane 8). Taken together, these results strongly support a mechanistic model whereby Hfq-dependent annealing of the PinT sRNA to this mRNA blocks the synthesis of the SPI-2 encoded effector SteC.

### PinT suppresses SteC-mediated actin rearrangement in host cells

When tested in standard *in-vitro* culture, deletion of *pinT* leads to only a marginal increase in SteC protein abundance (Fig. 3b), which is typical for sRNA deletion mutants (Hör et al., 2020). To test *steC* regulation in a more physiological setting, we infected HeLa cells with the *Salmonella steC::*3xFLAG strain. In this setting, we achieved to detect PinT and SteC, both expressed at physiological levels from the chromosome of intracellular *Salmonella* (Fig. 5a). Intriguingly, while the SteC::3xFLAG protein was not detectable at 4 h p.i. when infecting with wild-type *Salmonella*, it was already detectable at this point when the *pinT* deletion strain was used. Moreover, the Δ*pinT* strain sustained higher SteC levels throughout the infection up to 20 hours. Thus, the PinT-mediated repression indirectly delays SteC secretion into the host cells.

**Figure 5.**
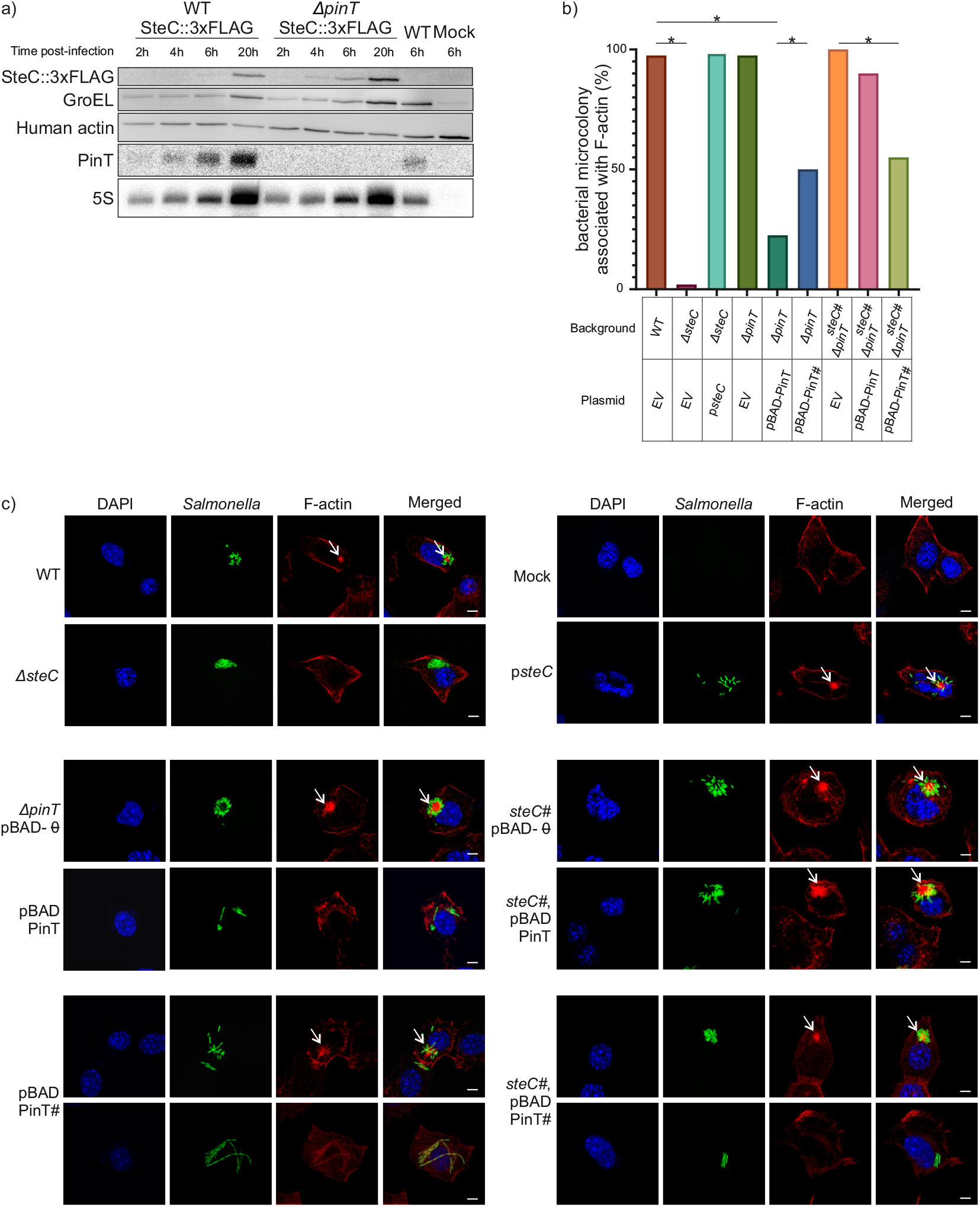
**a)** Time course expression of SteC. Western blot and Northern Blot analysis of intracellular *Salmonella* expressing SteC::3xFlag in the wild-type or a *pinT* deletion strains. Includes a wild-type *Salmonella* strain expressing wild-type SteC (non-tagged SteC) and a Mock control. HeLacells were infected with an M.O.I of 50. Protein and RNA samples were collected at 2h, 4h, 6h and 20h post infection. Includes GroEL, Human α-actin and 5S as a loading control for protein and RNA, respectively. Results from one out of two biological replicates are shown. **b)** Quantification of F-actin remodeling by *Salmonella* strains. Swiss 3T3 cells were infected for 10 h (M.O.I 100) with a constitutively GFP-expressing wild-type *Salmonella*, *steC* mutant, *pint* mutant, SteC complementation (under a constitutive promoter) or PinT complementation (under an arabinose inducible promoter). Also included are *Salmonella* strains with mutated 5’UTR of *steC* (*steC#* mutant) in the presence of over expression of PinT from a plasmid with an arabinose inducible promotor. Induction of PinT expression was achieved by adding arabinose directly to the media 1h post infection. Values correspond to the percentage of infected cells where F-actin was associated with the bacterial microcolony, at 10h post-infection. 40 infected cells for each strain were analyzed. Asterisk denotes significant differences between different strains (P<0.05; Fishers exact test). **c)** Representative confocal images of cells infected with GFP expressing *Salmonella* (green in merged image). F-actin was visualized by immuno-fluorescent labelling with Alexa Fluor 594 Phalloidin (red in merged image). Chromosomal DNA, shown in blue, was stained with DAPI.

The role of SteC is best understood in non-phagocytic cells where, following its translocation from *Salmonella*, this protein primarily functions as a kinase phosphorylating a specific set of cytosolic proteins involved in host immune signaling cascades that converge at the level of actin rearrangement (Imami et al., 2013; Odendall et al., 2012; Poh et al., 2008; Walch et al., 2020). Host protein phosphorylation by SteC is thought to trigger the formation of actin bundles in the vicinity of *Salmonella* microcolonies inside host cells. Therefore, actin arrangement lent itself as a host readout for determining the potency of PinT-mediated SteC repression during infection. To this end, we performed confocal microscopy to follow actin rearrangement in Swiss 3T3 fibroblasts, in which the SPI-2-dependent F-actin phenotype is particularly well defined (Méresse et al., 2001), infected with wild-type *Salmonella*, Δ*steC* or Δ*pinT* deletion mutants, and the corresponding complementation strains expressing either *steC* or *pin T* from a pZE-*luc* plasmid or a pBAD plasmid, respectively. In addition, all strains constitutively expressed GFP to track the localization of intracellular bacteria. Infected fibroblasts were fixed 10 h p.i., and actin filaments stained with an Alexa Fluor 595 phalloidin conjugate (Fig. 5c). Uninfected bystander cells (without a GFP signal) contained very few organized actin filaments (Fig. 5c). In agreement with previous studies (Poh et al., 2008), wild-type *Salmonella* caused the appearance of large clusters of condensed F-actin around the bacterial microcolonies in 97.5% of the infected cells. In case of the Δ*steC* mutant, this proportion dropped to 2% (Fig. 5b-c). *Trans*-complementation of *steC* in the Δ*steC* background restored actin rearrangement to 90% of the infected cells.

An effect of PinT on actin rearrangement was detectable when the sRNA was overexpressed. That is, while fibroblasts infected with Δ*pinT Salmonella* displayed the same numbers of infected cells with F-actin rearrangement as for the wild-type infection (97.5%), PinT expressed from an inducible promotor on a plasmid reduced this to 22.5%. This latter reduction suggested, but did not prove, that the PinT effect on actin formation was through repression of SteC synthesis.

Next, we screened actin rearrangement in fibroblasts infected with *Salmonella* engineered to carry a *steC*# allele on the chromosome (note that this allele is refractory to PinT activity; Fig. 3c-d). 100% of cells infected with this *Salmonella steC*# strain presented the F-actin phenotype; in other words, the *steC*# allele behaves like the wild-type *steC* gene. Interestingly, even when PinT was overexpressed, expression of *steC*# still resulted in actin rearrangement in 90% of the infected cells. However, the proportion of cells numbers with actin rearrangement dropped to 50% when the same *Salmonella* strain expressed PinT#, which harbors compensatory mutations for the *steC*# sequence. Interestingly, this reduction was similar when PinT# was expressed in a strain with a wild-type *steC* gene, which echoes the above results with *gfp* reporters (Fig. 3d). That is, the mutations present in PinT# allow for similar repression of the *steC* and *steC#* alleles. Nonetheless, the combined results support the notion that PinT, via repression of the *steC* mRNA, engages in the regulation of host actin organization during *Salmonella* infection.

## DISCUSSION

Recent methodological advances enabling high-resolution transcriptome studies in the context of host infection (Hör et al., 2018; Perez-Sepulveda and Hinton, 2018; Saliba et al., 2017; Westermann et al., 2017) have identified a plethora of sRNA genes that are co-regulated with virulence genes in bacteria (Caldelari et al., 2013; Quereda and Cossart, 2017; Svensson and Sharma, 2016; Westermann, 2018). Understanding the functions of these sRNAs in bacterial pathogenesis has been much harder: i) due to their short length, sRNA genes are often insufficiently covered in genomic screens for virulence factors, and ii) when investigated more systematically, sRNA deletions are rarely found to produce robust phenotypes in classical infection assays. Therefore, to understand how sRNAs contribute to the success of bacterial pathogens, a comprehensive bottom-up analysis of their molecular targets may be required. Our present findings with PinT argue that it is indeed worthwhile using different complementary methods in this endeavor.

Our MAPS-based discovery of the *steC* mRNA as a new PinT target naturally prompts the question of why it was overlooked in the two previous target searches for this sRNA. There reasons are different. One of the studies made use of educated guesses, selecting candidates with expected meaningful regulation around the invasion (SPI-1) phase of the *Salmonella* infection cycle, i.e., before SteC activity is relevant (Kim et al., 2019). The other study (Westermann et al., 2016) used sRNA pulse-expression, which in general terms has been very successful at predicting interactors of *E. coli* and *Salmonella* sRNAs (Hör et al., 2020). In that study, PinT was pulse-expressed for 5 min in different media as well as in *Salmonella* replicating inside epithelial cells, which was sufficient to downregulate other targets, i.e., *sopE* or *sopE2* (Westermann et al., 2016). Intriguingly, renewed inspection of that RNA-seq data shows that *steC* was downregulated, too, but did not pass as significant in any of the conditions tested. By contrast, MAPS with MS2-PinT clearly recovers *steC* as one of the top transcripts in two of the three conditions used here (Fig. 2). Particularly noteworthy is the SPI-2 MAPS experiment, which expresses the MS2-tagged sRNA from the native *pinT* locus to physiological levels. The successful enrichment of *steC* and other proven PinT targets suggests potential for reducing false-positives and dropouts in MAPS experiments, by avoiding overexpression of the fusion sRNA.

We have also attempted MAPS for *Salmonella* 4 hours into infection of macrophages, but the results are inconclusive such that most known PinT targets do not rank amongst the top. In our enrichment analysis, *grxA*, *sopE*, *steC*, and *sopE2* occupy ranks #222, #316, #980, and #1,226, out of listed 9878 features. While the possibility remains that the macrophage experiment enriches targets whose regulation is particularly relevant as *Salmonella* resides in this particular eukaryotic cell type and time of infection, these numbers rather suggest that successful intercellular MAPS is yet to be achieved. Improvement might come from a more rigorous depletion of contaminating eukaryotic RNA in the lysate; from the inclusion of an RNA-RNA cross-linking step; or from switching to other aptamers such as PP7 (Lim and Peabody, 2002; Tien et al., 2018) or Csy4 (Haurwitz et al., 2010; Lee et al., 2013) as sRNA fusion partners. Another possibility to increase sensitivity might be to combine MAPS with features of recent global RNA:RNA interactome techniques such as CLASH, GRIL-seq, or RIL-seq (Han et al., 2017; Melamed et al., 2018, 2016; Waters et al., 2017), e.g., purifying targets by stepwise pulldown of Hfq and the MS2-tagged sRNA.

What are the biological implications of the PinT-mediated repression of *steC* during infection? To promote intracellular replication, *Salmonella* remodels the host cytoskeleton and redirects a dense meshwork of F-actin in the vicinity of the SCV, and this process crucially hinges upon SteC (Odendall et al., 2012; Poh et al., 2008). Using different PinT and *steC* alleles, we have confirmed that PinT can regulate actin rearrangement in infected fibroblasts through repression of SteC protein synthesis (Fig. 5b-c). While these experiments required overexpression of PinT, we also provide evidence that for the effect of PinT on SteC levels under physiological conditions (Fig. 5a). SteC protein is first detected at 4 h p.i. (Poh et al., 2008) and its levels increase over time until 20 h p.i., while PinT levels peak at 8 h (Fig. 5a) (Westermann et al., 2016). We interpret this expression profile to mean that PinT functions to delay the arrival of SteC in the host cytosol until *Salmonella* has properly adapted its metabolism to the SCV environment. Moreover, SteC has a mild suppressive effect on bacterial growth (Poh et al., 2008), therefore delayed expression of this effector may be necessary to get intracellular bacterial replication going. The complex regulatory network, with the PhoP-induced sRNA PinT repressing PhoP-activated SteC, both directly and indirectly, make the *steC* mRNA a particularly interesting PinT target (Fig. 6), encouraging future studies of the relevance of these individual PinT-SteC-mediated changes to the host on a whole-organism level and for the outcome of infection. Such studies should also address how this repression may be counteracted, for example, through sequestration of PinT by other targets or by sRNA sponges, of which there is a growing number in bacteria (Figueroa-Bossi and Bossi, 2019).

**Figure 6.**
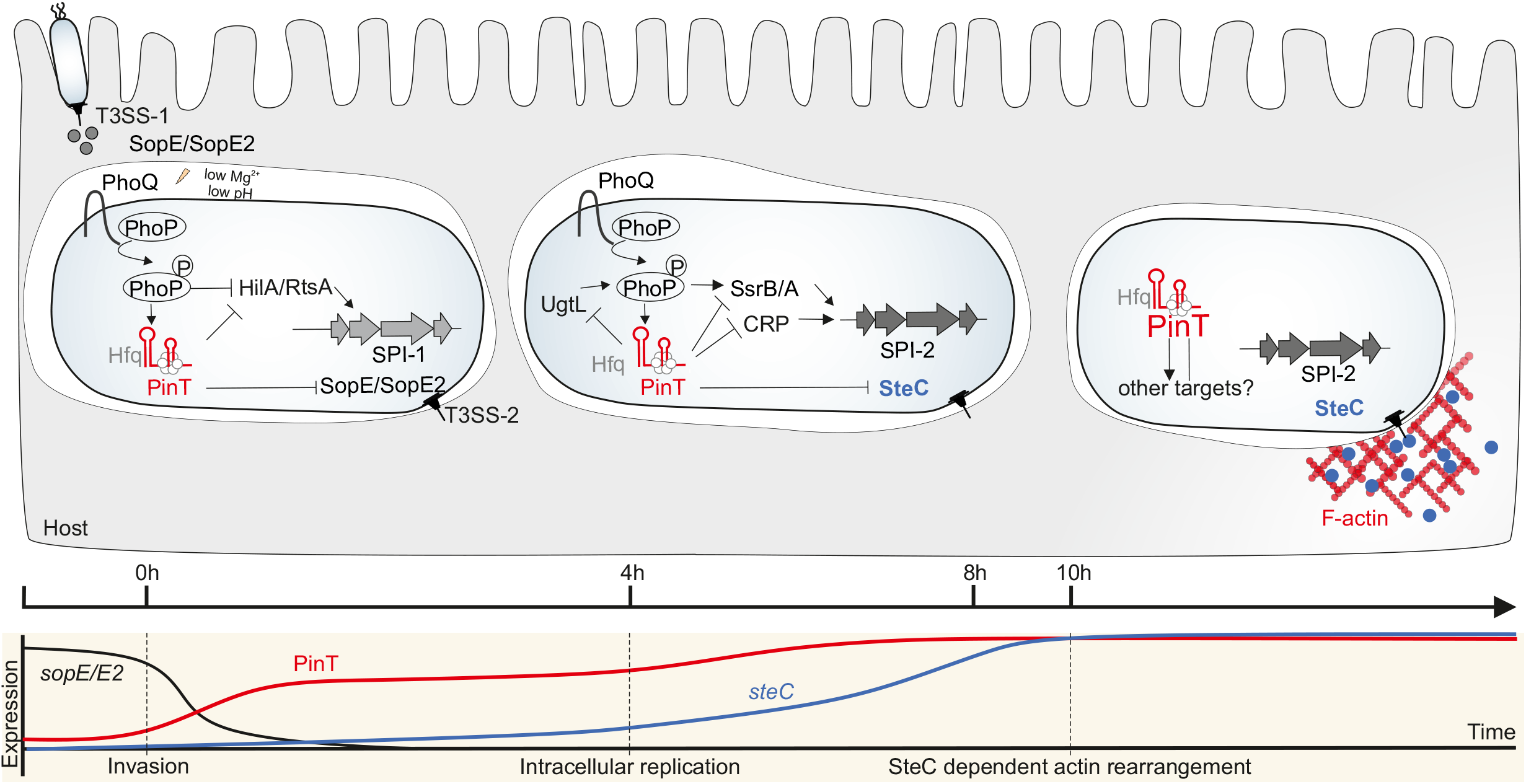
Current model of gene regulatory network of PinT during infection. Immediately after epithelial cell invasion, PinT expression is highly upregulated. At this time, expression of SteC is delayed by PinT binding to the *steC, crp* and *ssrB* mRNAs. Later in infection, repression by PinT is alleviated, by an unknown mechanism, and SteC is fully expressed. SteC production induces assembly of F-actin meshwork around the SCV. PinT provides an additional layer of crosstalk regulation between SPI-1 and SPI-2 virulence programs. This regulation occurs at several layers, with PinT post-transcriptionally repressing both SPI-1 and SPI-2 regulators, as well as SPI-1 and SPI-2 effector proteins. PhoQ, which activates PinT transcription and thus initiates the post-transcriptional control network schemed above, acts as a transcriptional SPI-1 repressor itself (Kim et al., 2019).

Finally, our MAPS datasets suggest that PinT may interact with even more RNA molecules than the ones currently validated. The mRNAs for the PhoQ activator UgtL and the 30S ribosomal protein RpsV were identified in all the different pull-downs. Interestingly, both *rpsV* and *ugtL* are highly upregulated in SPI-2 inducing conditions (Kröger et al., 2013) as well as inside macrophages (Srikumar et al., 2015). While we know little about the ribosomal protein RpsV, the *Salmonella*-specific inner membrane protein UgtL is known to mediate resistance to antimicrobial peptides by indirect modification of lipid A (Shi et al., 2004). UgtL is also required for gut colonization in streptomycin-treated mice (Goto et al., 2017). Most recently, UgtL was described as an activator of the PhoP/Q two-component system, contributing to *Salmonella* virulence in mice (Choi and Groisman, 2017). In other words, PinT is likely to repress the synthesis of an activator of its own transcription activator, potentially creating a feedback loop within PhoP/Q-mediated activation of SPI-2 genes (Fig. 6). As such, the emerging extended target suite of PinT promises to add new examples to the growing list of intermixed regulatory loops composed of sRNAs and transcription factors, which have been studied primarily in quorum sensing, metabolic responses, stress management as well as the transition between motility and sessility (Beisel and Storz, 2011; Mandin et al., 2016; Mouali et al., 2018; Shimoni et al., 2007). It will be interesting to see how the individual regulatory loops PinT is part of affect the timing of gene expression as *Salmonella* bacteria transition through different extracellular and cellular environments.

## METHODS

### Constructing Salmonella mutant strains and plasmids

We fused the MS2 aptamer (two steams of 48 nt in length each) to the 5’ end of PinT. To ensure the addition of the extra nucleotides to the 5’of the sRNA did not interfere with overall secondary structure linker nucleotides were added between the aptamers and the sRNA. This was verified using *in silico* RNA secondary structure prediction via *mfold* (http://unafold.rna.albany.edu/?q=mfold). A version of the tagged RNA in which MS2 is fused to the 5’ of PinT, was introduced in the pBAD plasmid, rendering MS2-PinT expression arabinose inducible. To construct the MS2 control plasmid (pYC-310) which expresses a short MS2 tag RNA, pYC128 was reamplified using oligos JVO-13630/2 and self-ligated. All plasmids used throughout this study were introduced into *Salmonella* by electroporation as described (Pfeiffer et al., 2007).

To express endogenous MS2-PinT, we adapted the λ Red recombinase method for one-step inactivation of chromosomal genes (Datsenko and Wanner, 2000). In summary, the DNA fragment to be integrated (containing either MS2-PinT or MS2-Term fused to the kanamycin resistance cassette of pKD4) was generated via overlapping PCR: the two DNA fragments obtained by PCR with primer pair JVO-16440/1 or JVO - 16440/9 (respectively) on gDNA and primer pair JVO-0203/16442 on pKD4 were mixed in equimolar amounts and used as template for a PCR with primer pairs JVO-16443/4. The chromosomal MS2-PinT or MS2-Term were subsequently transferred (transduced) to strains JVS-1574, cured with pCP20, resulting in strain JVS-12103 and JVS-12247.

### MS2-PinT overexpression

To test the expression levels of MS2-PinT and PinT, the strains SCS001 (Supplementary Table 2) were grown in LB with 50 μg/mL ampicillin (diluted 1:100 from an overnight culture). At OD_600nm_ of 2.0, expression of the plasmid was induced by adding 0.1% of arabinose to the media. After 10 minutes, 4 OD of cells were collected and mixed with StopMix. Total RNA was extracted using Trizol. 5 μg of total RNA were analyzed by Northern blot to verify that the MS2-PinT construct is expressed at a level similar to the untagged PinT construct (see Supplementary Table 2 for oligos).

### Time-course overexpression of PinT

To measure *sopE* and *sopE2* turnover after PinT overexpression, a strain with arabinose-induced overexpression of MS2-PinT (SCS001) was grown overnight and diluted 1:100 in LB with 50 μg/mL ampicillin at 37°C with shaking. At OD_600nm_ of 2.0, 4OD of cells were harvested. Next, 0.1% of arabinose was added to the media in order to induce MS2-PinT induction. Samples were collected at 1, 2, 5 and 10 minutes after induction, mixed with StopMix and immediately frozen in liquid nitrogen. Total RNA was extracted using Trizol extraction. RNA samples were treated with DNase I (Fermentas) for 45 min at 37°C. DNase I was then removed using phenol-chloroform extraction and RNA was again precipitated. DNase treatment was controlled by PCR using the oligos JVO-1224/1225 (Supplementary Table 2). qRT-PCR was performed with the Power SYBR Green RNA-to-CT 1-Step kit (Applied Biosystems) according to the manufacturer’s instructions. Fold changes for *pinT, sopE* and *sopE2* were determined using the 2^(−ΔΔCt)^ method relative to time before induction and using *5S* as reference gene (see Supplementary Table 2 for oligos).

### GFP reporter assay

To measure the GFP intensity of *sopE::gfp* reporter strains overexpressing MS2-PinT or PinT, bacteria were grown in LB in presence of Ampicillin and Chloramphenicol until an OD_600nm_ of 2.0. *Salmonella* cells corresponding to 1 OD were pelleted and fixed with 4% PFA. GFP fluorescence intensity was quantified for 100 000 events by flow cytometry with the FACSCalibur instrument (BD Biosciences). Data were analyzed using the Cyflogic software (CyFlo).

### MS2 affinity purification and RNA-seq

*Salmonella* strains, expressing either MS2-PinT (pYC362), PinT (pYC5-34), MS2 (pYC310) and under the control of pBAD inducible plasmid were used. Cells were grown in LB supplemented with 50 μg/mL ampicillin (diluted 1:100 from an overnight culture grown). At OD_600nm_ of 2.0, overexpression of the different constructs was induced by the addition of 0.1% arabinose. After two minutes, a volume of 60OD of cell was harvested and chilled on ice for 5 minutes. Cells were centrifuged at 10000 g for 10 minutes and the pellets frozen in liquid nitrogen. After thawing on ice, the pellets were resuspended in 600 μl of chilled Buffer A (20 mM Tris pH8.0, 150 mM KCL, 1 mM of MgCl2, 1 mM DTT). A volume of 750 μl of glass beads was added to the cells and lysed using Retsch (10 min, 30 Hz) (adaptors were pre-chilled at −20°C). Next, lysate was cleared by centrifugation 10 min at 16000g at 4°C and the clear lysate was collected into a new tube.

While the lysate is being prepared, affinity purification columns were prepared at the 4°C room. ~70 μl of amylose (New England Biolabs #E8021S) were added to 2mL Bio-Spin disposable chromatography columns (BioRad #732-6008). Amylose beads were washed three times with 2ml of Buffer A. 1ml of Buffer A with 250 pmol of MS2-MBP coat protein was added to the closed column followed incubation with rotation. After 5 minutes, the column was open and the MS2-coat protein was to run through the column and collected in a tube. This incubation step was repeated one more time until the lysate was ready. At this point, the solution was allowed to run and the column was washed once with 1ml of Buffer A.

The clear lysate was subjected to affinity chromatography (all the following steps were performed at 4 °C). Lysate was added to the closed column and incubated for 5 minutes with rotation. After the incubation, the lysate run was collected and the incubation step was repeated. Next, the column is washed 8 times with 2ml of Buffer A. Bound RNA was eluted using 300μl of Elution Buffer (Buffer A + 15 mM maltose). This step was repeated one more time. Eluted RNA was extracted with phenol-chloroform (V/V) and precipitated by the addition of ethanol (2 vol) and ~15μg of glycogen (1μl of Glycoblue 15mg/ml). RNA samples were treated with DNase I (Fermentas) for 45 min at 37°C. DNase I was then removed using phenol-chloroform extraction and RNA was again precipitated.

### MAPS in infected cells

The *in-vitro* infections of iBMMs (M.O.I 50) was carried out following a previously published protocol (Westermann et al., 2016) with slight modifications. Briefly, two days before infection 2 × 10^5^ iBMMs /mL were seeded in 10 ml complete DMEM (T75 flask). A total of 4 flaks of each cell type were infected with each *Salmonella* strain. Overnight cultures of *Salmonella* carrying either PinT (pYC55), MS2-PinT (pSS31) or MS2 (pSS32), were diluted 1:100 in fresh LB medium and grown aerobically to an OD_600nm_ of 2. Bacterial cells were harvested by centrifugation (2 min at 12,000 r.p.m., room temperature) and resuspended in DMEM. In iBMMs infection, bacterial cells were opsonized with 10% mouse serum for 20 min, prior to addition to the cells. After 4h of infection, the mammalian cells where washed with ice cold PBS and lysed with a solution of PBS 0.1% Triton. The bacterial cells where separated from the host cells debris by centrifugation at 500g for 5 min. After, the supernatant containing the bacteria was centrifuged again at 10 000 g for 5 min, washed with PBS and frozen. After elution, the purified RNA was sequenced and mapped to the either the Mouse genome and to the *Salmonella* genome.

### cDNA library preparation and sequencing

The RNA samples were first fragmented using ultrasound (4 pulses of 30 s each at 4°C). Then, an oligonucleotide adapter was ligated to the 3’ end of the RNA molecules. First-strand cDNA synthesis was performed using M-MLV reverse transcriptase and the 3’ adapter as primer. The first-strand cDNA was purified and the 5’ Illumina TruSeq sequencing adapter was ligated to the 3’ end of the antisense cDNA. The resulting cDNA was PCR-amplified to about 10-20 ng/μl using a high fidelity DNA polymerase (11 cycles). The TruSeq barcode sequences which are part of the 5’ TruSeq sequencing adapter are included in Table 2. The cDNA was purified using the Agencourt AMPure XP kit (Beckman Coulter Genomics) and was analyzed by capillary electrophoresis. For Illumina NextSeq sequencing, the samples were pooled in approximately equimolar amounts. The cDNA pool in the size range of 200-550 bp was eluted from a preparative agarose gel. The cDNA pool was paired-end sequenced on an Illumina NextSeq 500 system using 2×50 bp read length.

### Data analysis

Read counts were normalized by total read count for rRNA, using a scaling factor for each library. Next, we compared the ratio between the normalized read of the MS2-PinT sample and the PinT sample. Potential base-pairing with PinT was analyzed for the top hits of the list by using RNA-RNA interaction prediction tools, like IntaRNA software (Busch et al., 2008). Between the list of most enriched transcripts in the MS2-PinT sample and the PinT sample, nine candidate targets were selected based on their calculated free binding energy to PinT, infection relevance and SalComMac expression profile (Kröger et al., 2012; Srikumar et al., 2015).

### Time course pulse expression of PinT under OD2 and SPI2 conditions

To access the mRNa levels of candidate targets in response to PinT expression, a time course experiment of an overexpression of PinT was performed. Strains expressing PinT or the empty arabinose inducible plasmid were grown overnight and diluted 1:100 in LB at 37°C with shaking until OD_600nm_ of 2.0. In parallel, the same strains were grown in SPI-2 inducing conditions until OD_600nm_ of 0.3. Samples were collected before addition of arabinose and after 5, 10 and 20 of induction, mixed with StopMix and immediately frozen in liquid nitrogen. Total RNA was extracted using Hot Phenol extraction. DNase treatment was controlled by PCR using the oligos JVO-1224/1225 (Supplementary Table 2).

qRT–PCR was performed with the Power SYBR Green RNA-to-CT 1-Step kit (Applied Biosystems) according to the manufacturer’s instructions. Fold changes for pinT, *sopE, sopE2* and *steC* were determined using the 2^−ΔΔCt)^ method relative to time before induction and using 5S as reference gene (Supplementary Table 2).

### Detection of protein levels of candidate target genes

In order to quantify protein levels of the candidate target genes, a Flag tag fusion was added at the C-terminal end of the corresponding mRNA. JVS-3013 was transformed with PCR product from pSUB11. Oligos carried extensions (between 36 and 40 nt) homologous to the last portion of the targeted gene (forward primer) and to a region downstream from the stop codon (reverse primer) (Supplementary Table 2). P22 lysates were prepared from positive clones and used to transduce YCS-034. Insertion was verified by PCR and compared with the wild type (for oligos see Supplementary Table 2).

### In vitro structure probing and gel mobility shift assays

DNA templates that contain the T7 promoter sequence for *in-vitro* transcription using the MEGAscript T7 Kit (Ambion) were generated by PCR. Oligos and DNA templates used to generate the individual T7 templates are listed in Supplementary Table 2.

Gel-shift assays were performed with ~0.04 pmol 5’-end labeled PinT (4 nM final concentration) and increasing amounts of unlabeled RNA in 10 μl reactions. After denaturation (1 min at 95 °C), labeled RNAs were cooled for 5 min on ice and 1 μg yeast RNA and 10x RNA structure buffer (Ambion) were added. Increasing concentrations of unlabeled RNA were added to final concentrations of 12 nM, 47nM, 94 nM, 375nM, 750 nM, 1500 nM, and 3000 nM. For gelshift assays with ^32P^labeled *steC* mRNA truncated variant (from TSS to +250nt in the CDS), ~0.04 pmol labeled *steC* were incubated increasing concentrations of unlabeled PinT to final concentrations of 375 nM, 750 nM, 1500 nM, 2000 nM and 3000 nM. After incubation for 1h at 37 °C, samples were immediately loaded after addition of 3 μl 5x native loading dye (0.5 x TBE, 50% (vol/vol) glycerol, 0.2% (wt/vol) xylene cyanol and 0.2% (wt/vol) bromophenol blue) to a native 6% (vol/vol) PAA gel. For gelshifts with Hfq, ~0.04 pmol 5’-end labeled PinT was incubated increasing concentrations of unlabeled *steC* to final concentrations of 12 nM, 94 nM, 375 nM, 750 nM, 1500 nM and 3000 nM. Upon addition of purified Hfq 250nM) or Hfq dilution buffer (control), samples were incubated at 37°C for 1 h. Gel electrophoresis was done in 0.5 x TBE buffer at 300 V. Afterwards, gels were dried and analyzed using a PhosphoImager (FLA-3000 Series, Fuji) and AIDA software (Raytest, Germany).

Secondary structure probing and mapping of RNA complexes were conducted on 5’-end-labeled RNA (~0.1 pmol) in 10-μL reactions. RNA was denatured for 1 min at 95°C and chilled on ice for 5 min, upon which 1 μg of yeast RNA and 10× structure buffer (0.1 M Tris at pH 7, 1 M KCl, 0.1 M MgCl2; Ambion) were added. The concentrations of unlabeled sRNA/mRNA leader added to the reactions are given in the figure legends. Following incubation for 1 h at 37°C, 2 μL of a fresh solution of lead(II) acetate (25 mM; Fluka), 2 μL of RNase T1 (0.01 U/μL; Ambion), or 2 μL of Shortcut RNase III (0.02 L/μL; NEB) were added and incubated for 45 seconds, 3, or 10 min at 37°C, respectively. RNase III cleavage reactions contained 1 mM DTT and 1.3 U of enzyme (New England Biolabs #M0245S), and were incubated for 6 min at 37°C. Reactions were stopped with 12 μL of loading buffer on ice. RNase T1 ladders were obtained by incubating labeled RNA (~0.2 pmol) in 1× sequencing buffer (Ambion) for 1 min at 95°C. Subsequently, 1 μL of RNase T1 (0.1 U/μL) was added, and incubation was continued for 5 min at 37°C. OH ladders were generated by 5 min of incubation of 0.2 pmol of labeled RNA in alkaline hydrolysis buffer (Ambion) at 95°C. Reactions were stopped with 12 μL of loading buffer. Samples were denatured for 3 min at 95°C prior to separation on 6% polyacrylamide/7 M urea sequencing gels in 1× TBE. Gels were dried and analyzed using a PhosphorImager (FLA-3000 Series; Fuji), and AIDA software.

---

**Supplementary Figure 1.**
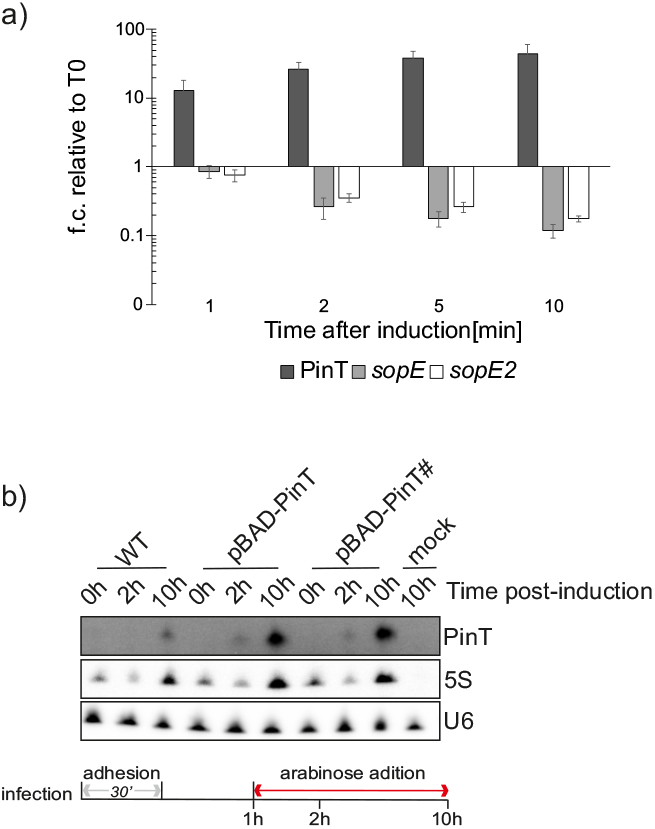
**a)** Time-course pulse expression of MS2-PinT. qRT-PCR measurements of PinT, *sopE* and *sopE2* over a time-course of 20 minute of MS2-PinT overexpression. Bars correspond to the mean fold change relative to before induction ± s.d. from three biological replicates and normalized to the control gene 5S. **b)** Northern Blot analysis of intracellular *Salmonella*. Swiss 3T3 cells were infected for 10 h (M.O.I 100) with a constitutively GFP-expressing wild-type *Salmonella* or the *pint* mutant, expressing either PinT or PinT# complementation under a arabinose inducible promoter. RNA samples were collected before addition of arabinose or 1h or 9h after induction. Includes 5S and U6 (mouse) as a loading control.

**Supplementary Figure 2.**
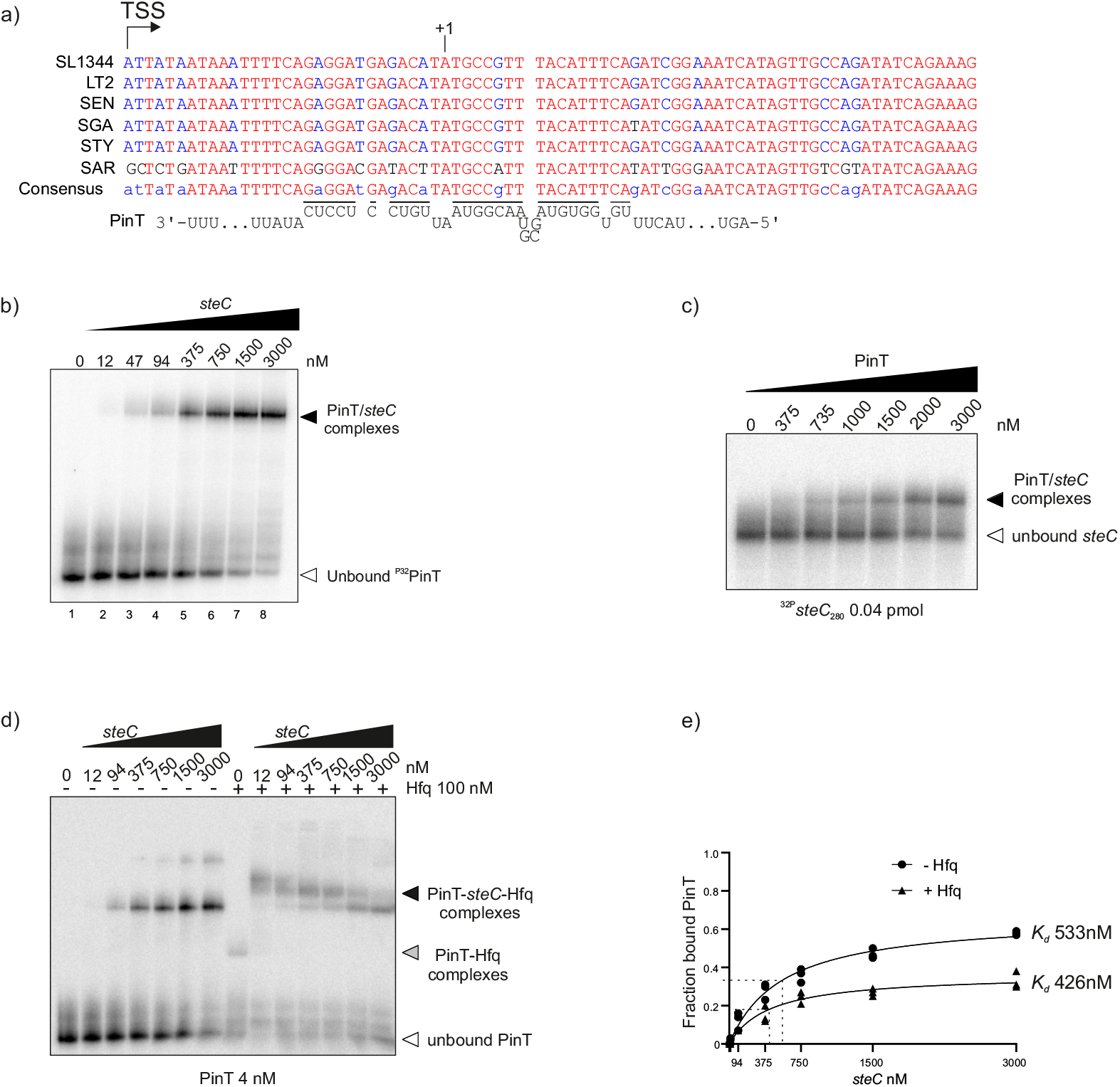
**a)** Sequence alignment showing the conservation of *steC* mRNA within the genus *Salmonella*. “STY”: S. Typhi, “SEN”: S. Enteritidis, “SGA”: S. Gallinarum, “SAR”: *S. arizonae*, “SBG”: *S. bongori*. Conserved ribonucleobases are labelled in red, less conserved bases are shown in blue. The numbers indicate the position relative to the start codon of *steC* (+1 position). Black lines and sequence motif below the alignment highlight the PinT binding site. **b)** Electrophoretic mobility shift assay (EMSA) of approximately 0.04 pmol ^32^P-labeled *steC* was incubated with increasing concentrations of unlabeled PinT. Full arrows indicate PinT/*steC* complex formation and empty arrows indicate unbound *steC*. **c)** Electrophoretic mobility shift assay (EMSA) of approximately 0.04 pmol ^32P^labeled PinT incubated with a fixed amount of Hfq and increasing concentrations of unlabeled *steC*. Full arrows indicate PinT/*steC* complex formation, grey arrows indicate PinT-Hfq complexes and empty arrows indicate unbound PinT. **d)** Quantitation of dissociation constant (Kd) values based on EMSA analysis from c).

## Acknowledgements

We are grateful to members of the Vogel and Westermann groups, to Lars Barquist and Jay Hinton for fruitful discussions of our research and constructive comments on the manuscript. We thank Elisa Venturini, Gianluca Matera, Daniel Ryan and Youssef El Mouali for comments on the manuscript. We thank the group of David Holden for the 3T3 Swiss fibroblasts and for guidance with the microscopy experiments. Thanks to Barbara Plaschke, Mona Alzheimer, Sarah Reichardt and Sandy Pernitzsch for technical support. This work was supported by funds from DFG (grant Vo875-19/1 and graduate program GRK 2157/1).

## REFERENCES

Ansong C, Yoon H, Porwollik S, Mottaz-Brewer H, Petritis BO, Jaitly N, Adkins JN, McClelland M, Heffron F, Smith RD. 2009. Global Systems-Level Analysis of Hfq and SmpB Deletion Mutants in Salmonella: Implications for Virulence and Global Protein Translation. PLoS One 4. doi:10.1371/journal.pone.0004809

Bardill JP, Zhao X, Hammer BK. 2011. The Vibrio cholerae quorum sensing response is mediated by Hfq-dependent sRNA/mRNA base pairing interactions. Molecular Microbiology 80:1381–1394. doi:10.1111/j.1365-2958.2011.07655.x

Barquist L, Westermann AJ, Vogel J. 2016. Molecular phenotyping of infection-associated small non-coding RNAs. Philos Trans R Soc Lond B Biol Sci 371. doi:10.1098/rstb.2016.0081

Beisel CL, Storz G. 2011. The Base-Pairing RNA Spot 42 Participates in a Multioutput Feedforward Loop to Help Enact Catabolite Repression in Escherichia coli. Molecular Cell 41:286–297. doi:10.1016/j.molcel.2010.12.027

Bobrovskyy M, Azam MS, Frandsen JK, Zhang J, Poddar A, Ma X, Henkin TM, Ha T, Vanderpool CK. 2019. Determinants of target prioritization and regulatory hierarchy for the bacterial small RNA SgrS. Molecular Microbiology 112:1199–1218. doi:10.1111/mmi.14355

Bradley ES, Bodi K, Ismail AM, Camilli A. 2011. A Genome-Wide Approach to Discovery of Small RNAs Involved in Regulation of Virulence in Vibrio cholerae. PLOS Pathogens 7:e1002126. doi:10.1371/journal.ppat.1002126

Busch A, Richter AS, Backofen R. 2008. IntaRNA: efficient prediction of bacterial sRNA targets incorporating target site accessibility and seed regions. Bioinformatics 24:2849–2856. doi:10.1093/bioinformatics/btn544

Cabezas CE, Briones AC, Aguirre C, Pardo-Esté C, Castro-Severyn J, Salinas CR, Baquedano MS, Hidalgo AA, Fuentes JA, Morales EH, Meneses CA, Castro-Nallar E, Saavedra CP. 2018. The transcription factor SlyA from Salmonella Typhimurium regulates genes in response to hydrogen peroxide and sodium hypochlorite. Research in Microbiology 169:263–278. doi:10.1016/j.resmic.2018.04.003

Caldelari I, Chao Y, Romby P, Vogel J. 2013. RNA-Mediated Regulation in Pathogenic Bacteria. Cold Spring Harb Perspect Med 3:a010298. doi:10.1101/cshperspect.a010298

Chao Y, Vogel J. 2010. The role of Hfq in bacterial pathogens. Current Opinion in Microbiology, Host-microbe interactions: bacteria 13:24–33. doi:10.1016/j.mib.2010.01.001

Chen J, Gottesman S. 2017. Hfq links translation repression to stress-induced mutagenesis in E. coli. Genes Dev 31:1382–1395. doi:10.1101/gad.302547.117

Choi J, Groisman EA. 2017. Activation of master virulence regulator PhoP in acidic pH requires the Salmonella-specific protein UgtL. Sci Signal 10:eaan6284. doi:10.1126/scisignal.aan6284

Colgan AM, Kröger C, Diard M, Hardt W-D, Puente JL, Sivasankaran SK, Hokamp K, Hinton JCD. 2016. The Impact of 18 Ancestral and Horizontally-Acquired Regulatory Proteins upon the Transcriptome and sRNA Landscape of Salmonella enterica serovar Typhimurium. PLOS Genetics 12:e1006258. doi:10.1371/journal.pgen.1006258

Datsenko KA, Wanner BL. 2000. One-step inactivation of chromosomal genes in Escherichia coli K-12 using PCR products. PNAS 97:6640–6645. doi:10.1073/pnas.120163297

Ellermeier JR, Slauch JM. 2007. Adaptation to the host environment: regulation of the SPI1 type III secretion system in Salmonella enterica serovar Typhimurium. Current Opinion in Microbiology, Host-microbe interactions: bacteria 10:24–29. doi:10.1016/j.mib.2006.12.002

Ellis MJ, Trussler RS, Charles O, Haniford DB. 2017. A transposon-derived small RNA regulates gene expression in Salmonella Typhimurium. Nucleic Acids Res 45: 5470–5486. doi:10.1093/nar/gkx094

Figueroa-Bossi N, Bossi L. 2019. Sponges and Predators in the Small RNA World. Regulating with RNA in Bacteria and Archaea 441–451. doi:10.1128/microbiolspec.RWR-0021-2018

Georg J, Lalaouna D, Hou S, Lott SC, Caldelari I, Marzi S, Hess WR, Romby P. 2020. The power of cooperation: Experimental and computational approaches in the functional characterization of bacterial sRNAs. Molecular Microbiology 113:603–612. doi:10.1111/mmi.14420

Gong H, Vu G-P, Bai Y, Chan E, Wu R, Yang E, Liu F, Lu S. 2011. A Salmonella Small Non-Coding RNA Facilitates Bacterial Invasion and Intracellular Replication by Modulating the Expression of Virulence Factors. PLOS Pathogens 7:e1002120. doi:10.1371/journal.ppat.1002120

Goto R, Miki T, Nakamura N, Fujimoto M, Okada N. 2017. Salmonella Typhimurium PagP- and UgtL-dependent resistance to antimicrobial peptides contributes to the gut colonization. PLOS ONE 12:e0190095. doi:10.1371/journal.pone.0190095

Guillet J, Hallier M, Felden B. 2013. Emerging Functions for the Staphylococcus aureus RNome. PLoS Pathog 9. doi:10.1371/journal.ppat.1003767

Han K, Tjaden B, Lory S. 2017. GRIL-seq provides a method for identifying direct targets of bacterial small regulatory RNA by in vivo proximity ligation. Nature Microbiology 2. doi:10.1038/nmicrobiol.2016.239

Hartz D, McPheeters DS, Traut R, Gold L. 1988. [27] Extension inhibition analysis of translation initiation complexesMethods in Enzymology, Ribosomes. Academic Press. pp. 419–425. doi:10.1016/S0076-6879(88)64058-4

Haurwitz RE, Jinek M, Wiedenheft B, Zhou K, Doudna JA. 2010. Sequence- and Structure-Specific RNA Processing by a CRISPR Endonuclease. Science 329:1355–1358. doi:10.1126/science.1192272

Hör J, Gorski SA, Vogel J. 2018. Bacterial RNA Biology on a Genome Scale. Molecular Cell 70:785–799. doi:10.1016/j.molcel.2017.12.023

Hör J, Matera G, Vogel J, Gottesman S, Storz G. 2020. Trans-Acting Small RNAs and Their Effects on Gene Expression in Escherichia coli and Salmonella enterica. EcoSal Plus 9. doi:10.1128/ecosalplus.ESP-0030-2019

Imami K, Bhavsar AP, Yu H, Brown NF, Rogers LD, Finlay BB, Foster LJ. 2013. Global Impact of Salmonella Pathogenicity Island 2-secreted Effectors on the Host Phosphoproteome. Molecular & Cellular Proteomics 12:1632–1643. doi:10.1074/mcp.M112.026161

Kim K, Palmer AD, Vanderpool CK, Slauch JM. 2019. The sRNA PinT contributes to PhoP-mediated regulation of the SPI1 T3SS in Salmonella enterica serovar Typhimurium. Journal of Bacteriology JB.00312-19. doi:10.1128/JB.00312-19

Kröger C, Colgan A, Srikumar S, Händler K, Sivasankaran SK, Hammarlöf DL, Canals R, Grissom JE, Conway T, Hokamp K, Hinton JCD. 2013. An Infection-Relevant Transcriptomic Compendium for Salmonella enterica Serovar Typhimurium. Cell Host & Microbe 14:683–695. doi:10.1016/j.chom.2013.11.010

Kröger C, Dillon SC, Cameron ADS, Papenfort K, Sivasankaran SK, Hokamp K, Chao Y, Sittka A, Hébrard M, Händler K, Colgan A, Leekitcharoenphon P, Langridge GC, Lohan AJ, Loftus B, Lucchini S, Ussery DW, Dorman CJ, Thomson NR, Vogel J, Hinton JCD. 2012. The transcriptional landscape and small RNAs of Salmonella enterica serovar Typhimurium. PNAS 109:E1277–E1286. doi:10.1073/pnas.1201061109

Lalaouna D, Baude J, Wu Z, Tomasini A, Chicher J, Marzi S, Vandenesch F, Romby P, Caldelari I, Moreau K. 2019a. RsaC sRNA modulates the oxidative stress response of Staphylococcus aureus during manganese starvation. Nucleic Acids Research 47:9871–9887. doi:10.1093/nar/gkz728

Lalaouna D, Carrier MC, Semsey S, Brouard JS, Wang J, Wade J, Massé E. 2015. A 3’ external transcribed spacer in a tRNA transcript acts as a sponge for small RNAs to prevent transcriptional noise. Molecular Cell 58:393–405. doi:10.1016/j.molcel.2015.03.013

Lalaouna D, Eyraud A, Devinck A, Prévost K, Massé E. 2019b. GcvB small RNA uses two distinct seed regions to regulate an extensive targetome. Molecular Microbiology 111:473–486. doi:10.1111/mmi.14168

Lalaouna D, Prévost K, Eyraud A, Massé E. 2017. Identification of unknown RNA partners using MAPS. Methods, RNA Methods in Bacteria 117:28–34. doi:10.1016/j.ymeth.2016.11.011

Lalaouna D, Prévost K, Laliberté G, Houé V, Massé E. 2018. Contrasting silencing mechanisms of the same target mRNA by two regulatory RNAs in Escherichia coli. Nucleic Acids Res 46:2600–2612. doi:10.1093/nar/gkx1287

Lee HY, Haurwitz RE, Apffel A, Zhou K, Smart B, Wenger CD, Laderman S, Bruhn L, Doudna JA. 2013. RNA–protein analysis using a conditional CRISPR nuclease. PNAS 110:5416–5421. doi:10.1073/pnas.1302807110

Lenz DH, Mok KC, Lilley BN, Kulkarni RV, Wingreen NS, Bassler BL. 2004. The Small RNA Chaperone Hfq and Multiple Small RNAs Control Quorum Sensing in Vibrio harveyi and Vibrio cholerae. Cell 118:69–82. doi:10.1016/j.cell.2004.06.009

Lim F, Peabody DS. 2002. RNA recognition site of PP7 coat protein. Nucleic Acids Res 30:4138–4144.

Löber S, Jäckel D, Kaiser N, Hensel M. 2006. Regulation of Salmonella pathogenicity island 2 genes by independent environmental signals. International Journal of Medical Microbiology 296:435–447. doi:10.1016/j.ijmm.2006.05.001

Mandin P, Chareyre S, Barras F. 2016. A Regulatory Circuit Composed of a Transcription Factor, IscR, and a Regulatory RNA, RyhB, Controls Fe-S Cluster Delivery. mBio 7:e00966–16, /mbio/7/5/e00966-16.atom. doi:10.1128/mBio.00966-16

Melamed S, Faigenbaum-Romm R, Peer A, Reiss N, Shechter O, Bar A, Altuvia Y, Argaman L, Margalit H. 2018. Mapping the small RNA interactome in bacteria using RIL-seq. Nat Protoc 13:1–33. doi:10.1038/nprot.2017.115

Melamed S, Peer A, Faigenbaum-Romm R, Gatt YE, Reiss N, Bar A, Altuvia Y, Argaman L, Margalit H. 2016. Global Mapping of Small RNA-Target Interactions in Bacteria. Molecular Cell 63:884–897. doi:10.1016/j.molcel.2016.07.026

Méresse S, Unsworth KE, Habermann A, Griffiths G, Fang F, Martínez-Lorenzo MJ, Waterman SR, Gorvel JP, Holden DW. 2001. Remodelling of the actin cytoskeleton is essential for replication of intravacuolar Salmonella. Cell Microbiol 3:567–577. doi:10.1046/j.1462-5822.2001.00141.x

Mouali YE, Gaviria-Cantin T, Sánchez-Romero MA, Gibert M, Westermann AJ, Vogel J, Balsalobre C. 2018. CRP-cAMP mediates silencing of Salmonella virulence at the post-transcriptional level. PLOS Genetics 14:e1007401. doi:10.1371/journal.pgen.1007401

Murphy ER, Payne SM. 2007. RyhB, an Iron-Responsive Small RNA Molecule, Regulates Shigella dysenteriae Virulence. Infection and Immunity 75:3470–3477. doi:10.1128/IAI.00112-07

Odendall C, Rolhion N, Förster A, Poh J, Lamont DJ, Liu M, Freemont PS, Catling AD, Holden DW. 2012. The Salmonella Kinase SteC Targets the MAP Kinase MEK to Regulate the Host Actin Cytoskeleton. Cell Host & Microbe 12:657–668. doi:10.1016/j.chom.2012.09.011

Padalon-Brauch G, Hershberg R, Elgrably-Weiss M, Baruch K, Rosenshine I, Margalit H, Altuvia S. 2008. Small RNAs encoded within genetic islands of Salmonella typhimurium show host-induced expression and role in virulence. Nucleic Acids Res 36:1913–1927. doi:10.1093/nar/gkn050

Papenfort K, Förstner KU, Cong J-P, Sharma CM, Bassler BL. 2015. Differential RNA-seq of Vibrio cholerae identifies the VqmR small RNA as a regulator of biofilm formation. PNAS 112:E766–E775. doi:10.1073/pnas.1500203112

Pérez-Morales D, Banda MM, Chau NYE, Salgado H, Martínez-Flores I, Ibarra JA, Ilyas B, Coombes BK, Bustamante VH. 2017. The transcriptional regulator SsrB is involved in a molecular switch controlling virulence lifestyles of Salmonella. PLOS Pathogens 13:e1006497. doi:10.1371/journal.ppat.1006497

Perez-Sepulveda BM, Hinton JCD. 2018. Functional Transcriptomics for Bacterial Gene Detectives. Microbiology Spectrum 6. doi:10.1128/microbiolspec.RWR-0033-2018

Pfeiffer V, Sittka A, Tomer R, Tedin K, Brinkmann V, Vogel J. 2007. A small non-coding RNA of the invasion gene island (SPI-1) represses outer membrane protein synthesis from the Salmonella core genome. Molecular Microbiology 66:1174–1191. doi:10.1111/j.1365-2958.2007.05991.x

Poh J, Odendall C, Spanos A, Boyle C, Liu M, Freemont P, Holden DW. 2008. SteC is a Salmonella kinase required for SPI-2-dependent F-actin remodelling. Cell Microbiol 10:20–30. doi:10.1111/j.1462-5822.2007.01010.x

Quereda JJ, Cossart P. 2017. Regulating Bacterial Virulence with RNA. Annual Review of Microbiology 71:263–280. doi:10.1146/annurev-micro-030117-020335

Said N, Rieder R, Hurwitz R, Deckert J, Urlaub H, Vogel J. 2009. In vivo expression and purification of aptamer-tagged small RNA regulators. Nucleic Acids Res 37:e133. doi:10.1093/nar/gkp719

Saliba A-E, C Santos S, Vogel J. 2017. New RNA-seq approaches for the study of bacterial pathogens. Current Opinion in Microbiology, Host-microbe interactions: bacteria 35:78–87. doi:10.1016/j.mib.2017.01.001

Shao Y, Bassler BL. 2014. Quorum regulatory small RNAs repress type VI secretion in Vibrio cholerae. Molecular Microbiology 92:921–930. doi:10.1111/mmi.12599

Shi Y, Cromie MJ, Hsu F-F, Turk J, Groisman EA. 2004. PhoP-regulated Salmonella resistance to the antimicrobial peptides magainin 2 and polymyxin B: Resistance to antimicrobial peptides. Molecular Microbiology 53:229–241. doi:10.1111/j.1365-2958.2004.04107.x

Shimoni Y, Friedlander G, Hetzroni G, Niv G, Altuvia S, Biham O, Margalit H. 2007. Regulation of gene expression by small non-coding RNAs: a quantitative view. Mol Syst Biol 3. doi:10.1038/msb4100181

Sievers S, Sternkopf Lillebæk EM, Jacobsen K, Lund A, Mollerup MS, Nielsen PK, Kallipolitis BH. 2014. A multicopy sRNA of *Listeria monocytogenes* regulates expression of the virulence adhesin LapB. Nucleic Acids Res 42:9383–9398. doi:10.1093/nar/gku630

Silva IJ, Barahona S, Eyraud A, Lalaouna D, Figueroa-Bossi N, Massé E, Arraiano CM. 2019. SraL sRNA interaction regulates the terminator by preventing premature transcription termination of rho mRNA. PNAS 116:3042–3051. doi:10.1073/pnas.1811589116

Sittka A, Pfeiffer V, Tedin K, Vogel J. 2007. The RNA chaperone Hfq is essential for the virulence of Salmonella typhimurium. Molecular Microbiology 63:193–217. doi:10.1111/j.1365-2958.2006.05489.x

Smith C, Stringer AM, Mao C, Palumbo MJ, Wade JT. 2016. Mapping the Regulatory Network for Salmonella enterica Serovar Typhimurium Invasion. mBio 7. doi:10.1128/mBio.01024-16

Sonnleitner E, Bläsi U. 2014. Regulation of Hfq by the RNA CrcZ in Pseudomonas aeruginosa Carbon Catabolite Repression. PLOS Genetics 10:e1004440. doi:10.1371/journal.pgen.1004440

Sonnleitner E, Gonzalez N, Sorger-Domenigg T, Heeb S, Richter AS, Backofen R, Williams P, Hüttenhofer A, Haas D, Bläsi U. 2011. The small RNA PhrS stimulates synthesis of the Pseudomonas aeruginosa quinolone signal. Molecular Microbiology 80:868–885. doi:10.1111/j.1365-2958.2011.07620.x

Srikumar S, Kröger C, Hébrard M, Colgan A, Owen SV, Sivasankaran SK, Cameron ADS, Hokamp K, Hinton JCD. 2015. RNA-seq Brings New Insights to the Intra-Macrophage Transcriptome of Salmonella Typhimurium. PLOS Pathogens 11:e1005262. doi:10.1371/journal.ppat.1005262

Svensson SL, Sharma CM. 2016. Small RNAs in Bacterial Virulence and Communication. Microbiology Spectrum 4. doi:10.1128/microbiolspec.VMBF-0028-2015

Tien M, Fiebig A, Crosson S. 2018. Gene network analysis identifies a central post-transcriptional regulator of cellular stress survival. eLife 7. doi:10.7554/eLife.33684

Tomasini A, Moreau K, Chicher J, Geissmann T, Vandenesch F, Romby P, Marzi S, Caldelari I. 2017. The RNA targetome of Staphylococcus aureus non-coding RNA RsaA: impact on cell surface properties and defense mechanisms. Nucleic Acids Res 45:6746–6760. doi:10.1093/nar/gkx219

Walch P, Selkrig J, Knodler LA, Rettel M, Stein F, Fernandez K, Viéitez C, Potel CM, Scholzen K, Geyer M, Rottner K, Steele-Mortimer O, Savitski MM, Holden DW, Typas A. 2020. Global mapping of Salmonella enterica-host protein-protein interactions during infection. bioRxiv 2020.05.04.075937. doi:10.1101/2020.05.04.075937

Waters SA, McAteer SP, Kudla G, Pang I, Deshpande NP, Amos TG, Leong KW, Wilkins MR, Strugnell R, Gally DL, Tollervey D, Tree JJ. 2017. Small RNA interactome of pathogenic E. coli revealed through crosslinking of RNase E. The EMBO Journal 36:374–387. doi:10.15252/embj.201694639

Westermann AJ. 2018. Regulatory RNAs in Virulence and Host-Microbe Interactions. Microbiol Spectr 6. doi:10.1128/microbiolspec.RWR-0002-2017

Westermann AJ, Barquist L, Vogel J. 2017. Resolving host–pathogen interactions by dual RNA-seq. PLOS Pathogens 13:e1006033. doi:10.1371/journal.ppat.1006033

Westermann AJ, Förstner KU, Amman F, Barquist L, Chao Y, Schulte LN, Müller L, Reinhardt R, Stadler PF, Vogel J. 2016. Dual RNA-seq unveils noncoding RNA functions in host–pathogen interactions. Nature 529:496–501. doi:10.1038/nature16547

Westermann AJ, Venturini E, Sellin ME, Förstner KU, Hardt W-D, Vogel J. 2019. The Major RNA-Binding Protein ProQ Impacts Virulence Gene Expression in Salmonella enterica Serovar Typhimurium. mBio 10. doi:10.1128/mBio.02504-18

